# Predicting binding affinity changes from long-distance mutations using MD simulations and Rosetta

**DOI:** 10.1101/2022.04.26.489494

**Authors:** Nicholas G. M. Wells, Colin A. Smith

## Abstract

Computationally modeling how mutations affect protein-protein binding not only helps uncover the biophysics of protein interfaces, but also enables the redesign and optimization of protein interactions. Traditional high-throughput methods for estimating binding free energy changes are currently limited to mutations directly at the interface due to difficulties in accurately modeling how long-distance mutations propagate their effects through the protein structure. However, the modeling and design of such mutations is of substantial interest as it allows for greater control and flexibility in protein design applications. We have developed a method that combines high-throughput Rosetta-based side-chain optimization with conformational sampling using classical molecular dynamics simulations, finding significant improvements in our ability to accurately predict long-distance mutational perturbations to protein binding. Our approach uses an analytical framework grounded in alchemical free energy calculations while enabling exploration of a vastly larger sequence space. When comparing to experimental data, we find that our method can predict internal long-distance mutational perturbations with a level of accuracy similar to that of traditional methods in predicting the effects of mutations at the protein-protein interface. This work represents a new and generalizable approach to optimize protein free energy landscapes for desired biological functions.

**Author Summary:** Protein-protein interactions are vital to almost all biological processes, and therefore the ability to accurately and efficiently predict how mutations alter protein binding has far-reaching applications in protein analysis and design. Current approaches to predict such mutational free energy changes are limited to mutations directly at the interaction interface. Much research has underlined the prevalence of allosteric protein regulation in biological processes, indicating the importance of understanding and predicting the effects of protein perturbations which act over long distances. In this work we develop a novel method based on molecular dynamics simulations, the Rosetta macromolecular modeling suite, and an analytical framework from alchemical free energy calculations which can predict the effects of long-distance mutations with levels of accuracy rivaling state of the art interface-specific methods. We hope that our method will serve as a novel framework for high throughput mutational analysis and therefore benefit future protein design efforts.

## Introduction

Protein interactions are central to many vital biological processes such as antibody binding, signal transduction, gene expression, and enzyme regulation. Consequently, changes in the propensity of proteins to associate or dissociate, for example due to mutation or post-translational modification, can be very disruptive to biological function and are therefore often linked to disease(1–3). The ability to understand and control the energetics of protein interactions therefore has far-reaching consequences for the study of disease and for the development of novel protein-based therapeutics.

A number of strategies have been developed to quantitatively predict the effects of mutations on protein interactions based on either empirical(4–6) or statistically-based(7, 8) scoring approaches, where energy functions or observations from known protein structures, respectively, are used to describe the physical interactions between protein residues and estimate free energy changes upon mutation. The relatively low computational cost of these scoring methods makes them well suited to large-scale analyses and protein design, albeit at the expense of accuracy(9). Modest improvements have been made to score-based approaches by integrating information from MD simulations to account for solvent effects and variations in protein conformation(10–12). Additionally, approaches which utilize the more physics-based scoring capabilities of the Rosetta macromolecular modeling suite have recently shown a promisingly high degree of accuracy in predicting mutational binding free energy changes(13–15), finding notable success in design applications(16, 17). Recently, machine learning algorithms and artificial intelligence have also been applied to score- based approaches, improving the prediction of interface-adjacent mutations(7, 18, 19). However, despite this progress, current scoring methods are almost exclusively limited to predicting the energetic effects of mutations at the protein interaction interface and fail to accurately predict the effects of mutations whose perturbations act over long distances.

Other approaches to calculating mutational free energy changes involve applying theoretically rigorous statistical mechanics and conformational sampling from MD simulations to calculate, rather than approximate through heuristics, mutational free energy changes(20–22). Free Energy Perturbation (FEP) is a method to determine the free energy difference between two states by simulating the system in one state while simultaneously evaluating the energies of the system in another state(22, 23). FEP is based on the Zwanzig equation(23), which states that the free energy difference between two states is equal to the exponential average of the energy differences between the two states:

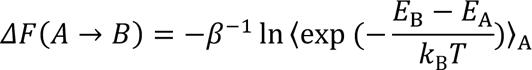

Here, *ΔF*(*A → B*) is the free energy difference between states *A* and *B*, β is equal to the inverse of the product of temperature (*T*) and Boltzmann’s constant (*k*_B_), *E*_A_ and *E*_B_ are the energies of conformations in state A and B respectively, and angular brackets denote the average over conformations taken from an equilibrium ensemble of state A.

While this theorem is numerically exact with an infinite sample of energy differences and only requires simulation of one equilibrium ensemble, it is highly reliant on sufficient sampling of the equilibrium system to achieve acceptable levels of accuracy and convergence(24, 25). It also often requires long and computationally expensive MD simulations simulated at many intermediate states when applied to typical biomolecular systems.

Methods like the Bennet acceptance ratio (BAR) and thermodynamic integration (TI) are non-equilibrium alternatives to FEP which involve calculating the work of a transition in the “forward” as well as the “reverse” directions, i.e. 6_A→B_ and 6_B→A_(26).

Such methods have shown high degrees of accuracy in predicting the effects of mutations, both interface-adjacent and allosteric, on biophysical processes such as protein association(27, 28). However, while these “bidirectional” approaches significantly improve the accuracy and convergence of free energy estimates compared to their “unidirectional” counterparts(29), they further increase the need for computationally expensive MD simulations by requiring separate equilibrium simulations for each state of interest to allow calculation of a set of “reverse” work values. In addition to substantial equilibrium sampling, these methods also often rely on computationally expensive simulations to generate non-equilibrium work values(20, 27, 29), making them less suitable for large-scale applications.

Recent studies have emphasized the importance of allosteric regulation in protein interactions throughout biology(30–32), underscoring the power of long-distance perturbations to protein structure and function. The ability to predict the effects of long-distance mutations would significantly improve the current scope of protein design and allow for a greater understanding of disease processes which feature long-distance protein modulation. To the best of our knowledge there does not currently exist an accurate high-throughput method to predict the effects of long-distance mutations on protein-protein interactions. In the present study, we develop the Molecular Dynamics Rosetta-Free Energy Perturbation (MDR-FEP) method which combines the extensive conformational sampling from classical MD simulations with the high-throughput nature of Rosetta repacking and the theoretical framework of alchemical FEP, allowing for the accurate prediction of long-distance mutational free energy changes with a high enough throughput for large-scale mutational analyses and forward protein design.

## Results and Discussion

### Protocol Rationale and Overview

The effect of mutations directly at the protein-protein interface can often be modeled using algorithms that keep the backbone fixed and allow side chain flexibility(15). However, the farther a mutation is away from the protein-protein interface, the more likely it is to affect binding through backbone perturbations like shifts in secondary structure or protein loops. Prediction of those backbone shifts can be done with two different strategies. An “induced-fit” approach involves first making the mutation and then using structural refinement algorithms to predict how the mutation propagates changes to the rest of the structure and alters the binding free energy. Alternatively, a “conformational selection” approach involves first obtaining an ensemble of backbone structures with the wild-type sequence and then later determining whether given mutations will sufficiently alter the energetics of that backbone ensemble to affect the binding free energy.

For the purposes of this work, we assume that sampling of backbone conformations is significantly more computationally costly than sampling of side chain conformations. This is largely supported by the high performance of algorithms like the Rosetta packer(33), which can very efficiently sample both rotamer and sequence space, and the considerably more complex task of backbone sampling. Given this dramatic difference in performance between the sampling of backbone and side chain conformations, a “conformational selection” algorithm that creates an ensemble of backbone conformations once and then samples sequence space many times is likely to be much more efficient than an “induced-fit” approach. While a “conformational selection” algorithm may perform poorly on mutations that require larger shifts of the backbone ensemble than would be captured in an “induced-fit” approach, there likely remains a large number of mutations that represent low hanging fruit amenable to conformational selection.

We hypothesized that equilibrium molecular dynamics simulations of a WT protein would serve as an effective means of generating a backbone ensemble for a “conformational selection” approach. To be effective, the MD simulations should transiently sample backbone conformations for which the energy with a given mutant sidechain is lower than WT, providing information about that mutation’s relative ΔG in the free or bound state. Because backbone conformations which are more stable with a mutant sidechain could have higher energies than with the WT sidechain present, these conformations may be infrequently sampled in WT simulations. Previous studies(12, 34, 35) found improved accuracy in predicting the effects of interface-adjacent mutations using the FoldX method after averaging values from structural snapshots of an MD simulation as compared to using only a single structure. However, due to the increased difficulty of predicting long-distance mutations, we anticipated that the WT simulations would sample long-distance mutation-accommodating conformations less frequently.

Such infrequent sampling would lead to a relatively small contribution of these low-energy conformations to the overall ensemble, resulting in negligible shifts in the mean or median energies for the WT and mutant sequences. To detect such shifts at the extremes of the energy distributions, we tested whether the use of analytical methods rooted in statistical mechanics enables better accounting for the contribution of these low-energy conformations to the overall ΔG value. In the present work, the Rosetta packer is used to side-chain optimize and score the energy of each frame in an equilibrium simulation with both WT and mutant sequences. These energies are then evaluated using exponential averaging in the Zwanzig equation (above) to produce a ΔG value.

The β parameter of that function, which describes the inverse of the equilibrated system’s theoretical temperature multiplied by the Boltzmann constant, dictates the extent to which low-energy conformations contribute to ΔG: At low values of β, frames with a high ΔE dominate while those with a low ΔE contribute very little. Conversely, an increase in β causes the contribution of low-energy conformations to increase (Figure S1). While the MD simulations are run at a well-defined temperature with known β, the Rosetta packer uses simulated annealing to optimize the total energy over a set of discrete rotamer configurations. Packing is followed by gradient-based sidechain dihedral angle minimization. As a result of these distinctly non-equilibrium processes, the method is no longer theoretically rigorous like standard FEP and an appropriate temperature is not known *a priori*. Due to these theoretical and practical differences, an alternative variable name other than β may be preferable, but we continue to use it here to maintain continuity with the original Zwanzig equation. Many parts of the Rosetta energy function use functional forms grounded in physics whose parameters are empirically optimized using structural data. Likewise, in the proposed method, β is treated as a parameter to be optimized in order to find the ideal level of contribution of low-energy frames to ΔG.

For a protein dimer of chains A and B, WT MD simulations are run with both dimeric and monomeric chains (Figure 1A, Step 1). Individual structural snapshots are extracted every 1 ns from each trajectory. For each frame, all residues within 10 Å of the mutation of interest are repacked and scored with both the WT and mutant sidechain present, generating a distribution of ΔE values for both the monomer and dimer states (Figure 1A, Step 2). In order to account for the possibility of outliers, a cutoff is then applied to this density function where all values which fall below the cutoff are set equal to zero. The set of ΔE values generated for each mutation in the monomer and dimer states is converted into a distribution using kernel density estimation (KDE) in order to allow for the application of more precise cutoff values (Figure 1B-C). With no cutoff applied, MDR-FEP performance was comparable when using KDE and raw ΔE values (Figure S2). However, KDE allowed for the continuous exclusion of a percentage of the density curve allowing for more precise control while analysis of raw ΔE values required discretely excluding whole frames (Figure S3). The Zwanzig equation is then applied to each distribution to generate a ΔG value for the monomer and dimer, allowing for the calculation of ΔΔGDimerization for the mutation of interest.

**Figure 1.**
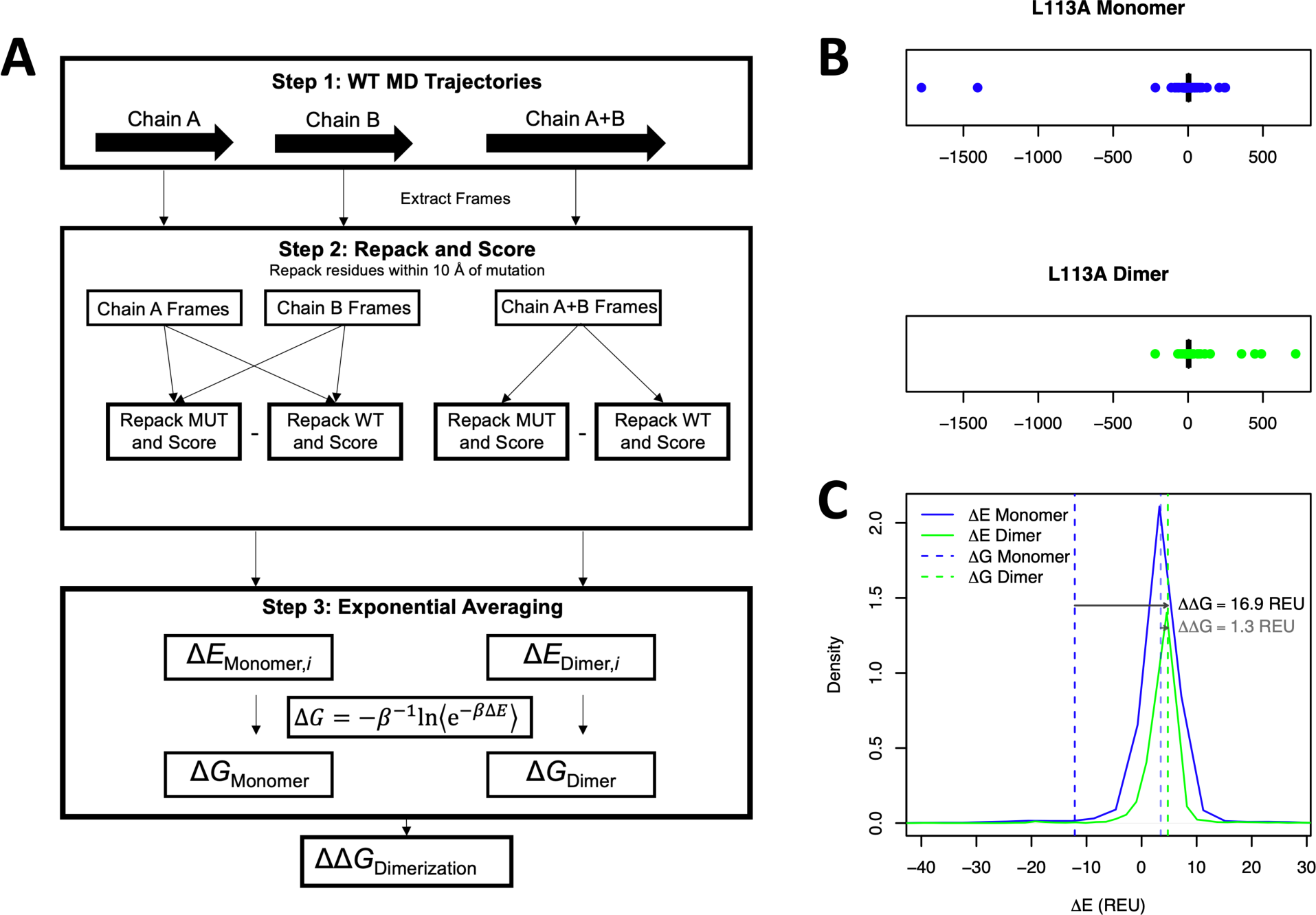
Schematic of the MDR-FEP method. **(A)** A schematic overview of the proposed method is shown, with the first step involving Gromacs molecular dynamics simulations, the second step using Rosetta repacking simulations, and the third step involving analysis of the resulting energies. **(B)** Box plots containing all ΔE values for the L113A monomer and dimer are shown, where each frame outside the interquartile range represented by a dot. **(C)** Images of the primary ΔE distributions for the L113A monomer and dimer are shown. ΔG values, calculated using β = 0.002 and cutoff = 0.005%, are shown as dashed lines using all frames (solid colors) and using only frames with ΔE > -500 REU (opaque colors). The final ΔΔG value is the difference between the blue and green dashed lines, represented by a solid arrow for the calculation including all frames and by an opaque arrow for the calculation using only frames with ΔE > -500 REU.

### Parameter Optimization and Performance Evaluation

We first performed a grid search over three separate mutation datasets in order to determine optimal β and cutoff parameters for the proposed method (Figure 2). To evaluate how well our method recapitulated experimental data at each given parameter set, Pearson correlation with experimental data was evaluated with β ranging from 0 to 0.1 and cutoff ranging between 0 and 1%. We first evaluated our method against an experimental surface plasmon resonance alanine scan(36) of urokinase-urokinase receptor dimerization for both interior long-distance and interface-adjacent mutations, which included experimentally validated values showing long-distance mutations which both stabilize and destabilize protein binding (Figure 2A-B). Interestingly, we found that the method’s performance in predicting the effects of both long-distance and interface mutations in the urokinase-urokinase receptor interaction seemed to depend on the cutoff parameter, with correlations appearing to decrease with cutoffs larger than 0.05% and 0.03%, respectively (Figure 2A/B). We interpret these results to indicate that the accurate prediction of the ΔG of these mutations relies on the sampling and correct accounting of low-energy frames, perhaps due to the increased difficulty of simulating long-distance and sterically larger mutations to alanine. To investigate this finding further, we evaluated MDR-FEP performance using simple averaging of ΔE distributions rather than exponential averaging, where low energy frames are able to shift the calculated value significantly less. As expected, MDR-FEP correlation to experiment for both long-range interior and interface mutations were lower and statistically insignificant (R = 0.03 and 0.13, respectively; p > 0.05) when using simple averaging (Figure S4A/B). Conversely, the correlation for each of these sets of mutations seemed to depend less on the β parameter (Figure 2A/B).

**Figure 2.**
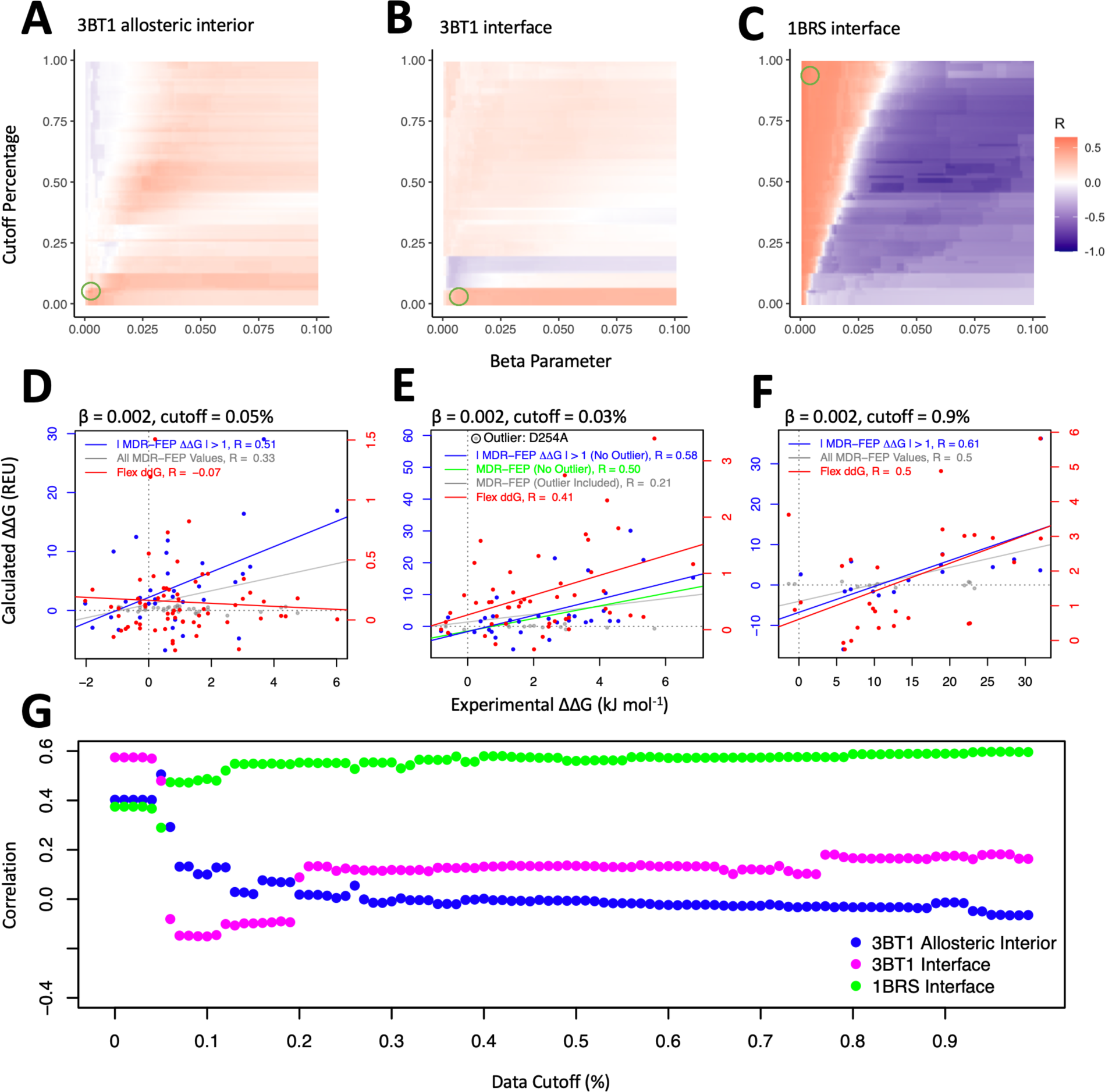
Parameter optimization by grid search. **(A-C)** Grid search heat maps showing the correlation of MDR-FEP predicted data to experiment for (A) 3BT1 allosteric, (B) 3BT1 interface, and (C) 1BRS interface sets of mutations. Correlations were calculated after excluding negligible predicted ΔΔG mutations for all datasets. Green circles indicate the set of parameters producing the highest correlation. **(D-F)** The performance of the MDR-FEP method is compared to Flex ddG(13), a leading Rosetta-based ΔΔG calculation method, for each set of mutations in (A-C) shown above. **(G)** The correlation of each set of mutations in (A-C) is plotted against the data cutoff using β = 0.002. Correlations were calculated after removal of negligible predicted ΔΔG mutations for all datasets, and after the removal of the outlier indicated by the gray circle in (E) for 3BT1 interface mutations.

We then evaluated the MDR-FEP method using an experimental ITC study(37) of the barnase-barstar dimerization process which included only non-allosteric (interface-adjacent) mutations (Figure 2C), finding that our method was able to reproduce experimental data with reasonable accuracy independent of the cutoff parameter while using a low β value. Correlation with the experimental ITC data did however decrease sharply at higher β values (Figure 2C). The fact that correlation did not decrease as more low-energy frames were removed by the increasing cutoff parameter suggests that the accurate prediction of ΔG for these mutations does not rely on the sampling of low-energy frames. Correlation with experiment remained approximately the same (R = 0.5, p = 0.03) using simple averaging (Figure S4C), further indicating that low-energy frames are not as important for this easier interface-adjacent mutation dataset. Although the 1BRS and 3BT1 simulations sampled approximately the same number of low-ΔG conformations (Figure S5), we believe that mutations in the 1BRS system are overall easier to predict by Rosetta repacking alone and therefore rely less on the sampling and accounting of frames which approximate mutation-accommodating backbone conformations.

We then compared the performance of the proposed method to that of Flex ddG(13), an existing Rosetta-based ΔΔGDimerization calculation method. Over a set of 65 allosteric mutations to alanine (Figure 2D), our method achieved an overall correlation of 0.33 (p = 7.4x10^-3^) while Flex ddG showed a correlation of approximately 0 (p = 5.9x10^-1^). While clearly an improvement over Flex ddG, the somewhat low MDR-FEP correlation coefficient was expected given the difficulty of predicting binding free energy changes using only a conformational selection approach at the backbone level. For certain positions and/or mutations, this approach is likely to fail due to rare sampling of the backbone conformations necessary for accurate predictions. We therefore sought to identify a simple heuristic that could be used to prospectively screen out mutations that are not likely to be predicted correctly. In the context of protein design, those mutations could then be excluded from experimental validation, increasing the likelihood of success at either stabilizing or destabilizing protein-protein binding. We noticed that a number of mutations were falsely predicted to have ΔΔG values between -1 and 1 REU by the MDR-FEP method. We term these “negligible predicted ΔΔG” mutations because the algorithm predicts a negligible ΔΔG (correctly or incorrectly), not because the actual ΔΔG is negligible. Interestingly, correlation improved significantly (R = 0.33 to 0.51; p = 7.4x10^-3^ to 2.6x10^-3^) with the exclusion of these “negligible predicted ΔΔG” mutations whereas Flex ddG did not show any improvement when applying this same filter. This improvement may result from the exclusion of mutations which feature extensive backbone alterations and are therefore too difficult for the MDR-FEP method to predict using the current level of backbone sampling.

Over a set of 49 mutations to alanine at the urokinase-urokinase receptor interface (Figure 2E), the proposed method achieved an overall correlation of 0.21 (p = 1.6x10^-1^) compared to Flex ddG’s 0.41 (p = 3.0x10^-3^). We observed that the MDR-FEP method was predicting very negative ΔG values for the D254A mutation in the monomer, even at low β values. Analysis of this mutation revealed that less than 90% of the contribution to the exponential averaging fell within the main distribution, while for all other mutations this was not the case. This analysis indicates an inappropriately large contribution from low-energy conformations for this particular mutation (Figure S6).

Exclusion of this potential outlier resulted in a correlation of 0.50 with experiment and increased statistical significance (p = 3.1x10^-4^). The correlation for the proposed method improved to 0.58 (p = 3.9x10^-4^) with the exclusion of negligible predicted ΔΔG mutations and the outlier, while the correlation of Flex ddG did not show such an improvement. Finally, for interface-adjacent barnase-barstar mutations, both the MDR-FEP method and Flex ddG achieved an overall correlation to experiment of approximately 0.5 (p = 6.3x10^-3^ and 7.1x10^-3^, respectively) over a set of 28 mutations (Figure 2F). Again, the MDR-FEP method showed an improved correlation with experiment (R = 0.5 to 0.61, p = 6.3x10^-3^ to 1.1x10^-2^) with the exclusion of negligible predicted ΔΔG mutations, while Flex ddG did not. We noticed that the MDR-FEP method was predicting larger-magnitude ΔΔG values for the W35F and H102L mutations which appeared to contribute significantly to the overall correlation. In order to evaluate performance without these potential outliers, we analyzed the correlation after excluding these mutations from the dataset. This analysis revealed a lower but still significant correlation to experiment (0.41 vs 0.50, p < 0.05) (Figure S7).

After observing that the optimal β value for all three datasets was approximately 0.002, we sought to determine an optimal consensus cutoff parameter that would maximize correlation independent of the dataset. We compared the correlation of each dataset using a β value of 0.002 against cutoff values ranging from 0 to 1% (Figure 2G), finding that all three datasets demonstrated acceptable correlations with no cutoff applied. Based on these results, we propose an optimal consensus parameter set of β =0.002 and cutoff = 0%.

The absolute scale of our ΔΔG predictions was significantly higher than that of the experimentally determined values. The REU scale is calibrated to be roughly comparable to kcal/mol(38), which further increases the discrepancy in overall scale with the experimental values shown in kJ/mol. However, that calibration may no longer be valid because of the Zwanzig algorithm’s emphasis on very low energy structures. The combination of the Rosetta packer and sidechain dihedral angle minimization may also exaggerate the magnitude of the energy differences found. Another factor that may exaggerate the absolute value and/or error of the predicted ΔΔG is differences in the ideal sidechain bond lengths and angles used by the AMBER 99SB*-ILDN force field and the Rosetta packer. The “idealization” of bond geometries during Rosetta rotamer generation may introduce clashes or unsatisfied hydrogen bonds that cannot be resolved by dihedral angle minimization alone. Investigation of these effects, especially in the context of other systems with experimentally characterized long-distance mutations, may lead to further improvements in the MDR-FEP algorithm.

### Predictive Performance Depends on Mutation Location

After observing good performance with both interface and long-distance mutations, we evaluated the performance of our method based on the location of the mutations included in the present study (Figure 3). Because the prediction of mutational effects becomes increasingly difficult at long distances, we hypothesized that ΔG values for 3BT1 mutations at extreme distances would be predicted with less accuracy. Indeed, we observed that mutations which were correctly predicted to be destabilizing or stabilizing fell closest to the 3BT1 interface (Figure 3A-C, green and blue). Likewise, mutations incorrectly predicted to be destabilizing or stabilizing tended to fall farther from the interface (Figure 3A-C, red and magenta). Interestingly, we observed no clear trend to explain the performance of negligible predicted ΔΔG mutations based on distance to the interface. This analysis may suggest that negligible predicted ΔΔG mutations may be harder to predict due to their more significant backbone alterations rather than their distance to the protein-protein interface. Additionally, we evaluated the performance of the MDR-FEP method at various distance cutoffs, where only mutations that fall closer to the interface than the cutoff are included in the calculation (Figure 3D). We found that correlation indeed decreased with the inclusion of more distant mutations.

**Figure 3.**
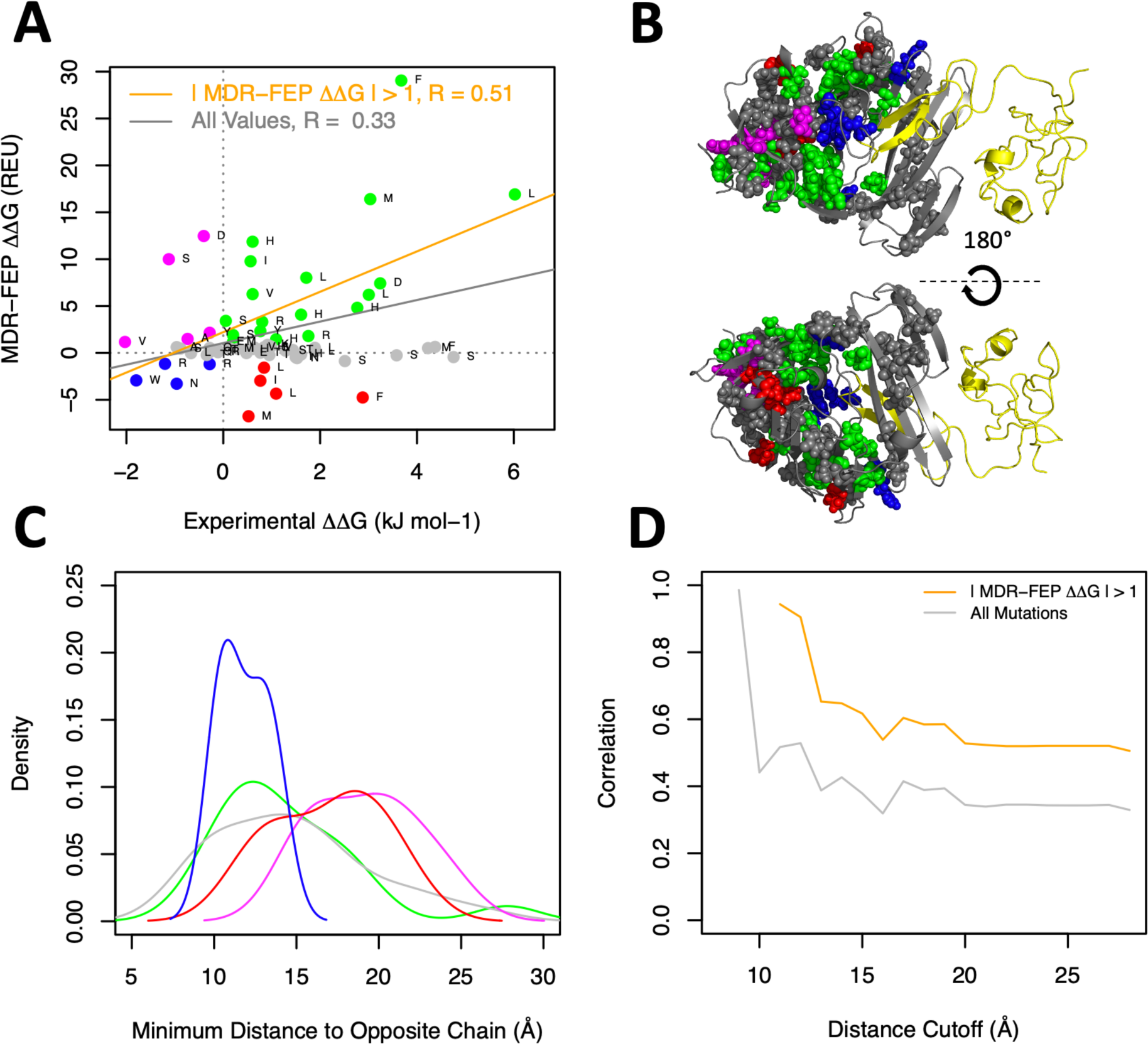
Calculated ΔΔGDimerization values show good agreement with experimental data for allosterically active mutations in the 3BT1 interior. **(A)** ΔΔGDimerization values calculated with the optimal parameter set (β = 0.002, cutoff= 0.05%) are plotted against experimentally determined values for 3BT1 mutations which are defined as “interior” by the SKEMPI database(47), meaning that these residues are located away from the protein-protein interface and have a relative solvent-accessible surface area of less than 25%. Lines of best fit are shown for the comparison of all mutations (gray) and for the set which excludes negligible predicted ΔΔG mutations (| MDR-FEP ΔΔG | ≤ 1 REU, orange). Dots are labeled by the identity of the residue being mutated to alanine (WT alanines are mutated to serine). Values which were correctly identified as stabilizing and destabilizing are shown in blue and green, respectively. Values which are incorrectly predicted to be stabilizing and destabilizing are shown in red and magenta, respectively. Negligible predicted ΔΔG mutations are shown in gray. **(B)** Mutations shown in Panel A are depicted as spheres on 3BT1 chain U (gray backbone) along with the binding partner chain A (yellow backbone). The sphere coloring corresponds to the dot coloring of Panel A. **(C)** Distributions of the minimum distance between a mutation’s Cβ and the closest Cβ (or Cα in the case of glycine) of the opposite chain for each set of mutations in (A). Colored based on (A-B). **(D)** MDR-FEP correlation with experiment, calculated using the same mutations and parameter set as in (A), is plotted against a distance cutoff where mutations with a minimum distance to the opposite chain according to (C) greater than the cutoff are excluded from the calculation. Colored according to (A).

We hypothesized that the exclusion of negligible predicted ΔΔG mutations improves correlation by removing “false negative” mutations, or mutations which are falsely predicted to have near-zero ΔΔG values. These false negatives likely arise due to a higher energy difference between the WT and mutant proteins, resulting in a smaller likelihood of sampling the low-energy frames needed to recapitulate experimental data. Indeed, we observed that negligible predicted ΔΔG mutations tend to sample fewer low-energy frames than their high predicted-value counterparts (Figure S8). Additionally, we found that 13 of the 36 negligible predicted ΔΔG allosteric mutations feature amino acids with hydroxyl groups (Figure 4A). Mutations which remove or introduce hydrogen bonds to the protein structure may be more difficult for the WT equilibrium simulations to sample, perhaps helping to explain this trend.

**Figure 4.**
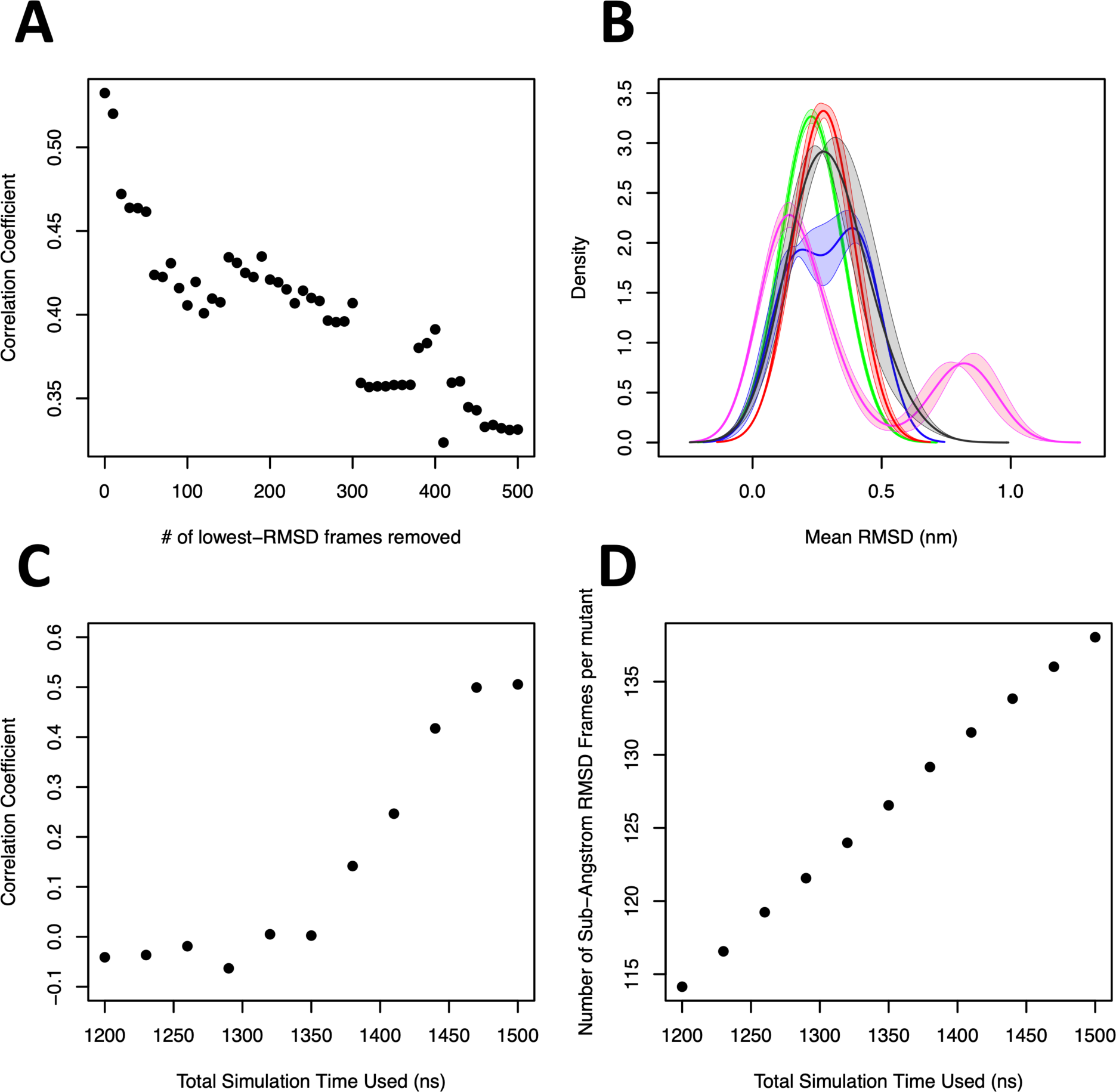
The MDR-FEP method performs well by sampling mutant-like conformations. **(A)** The correlation of predicted values to experimental data for all non-charge-changing interior allosteric 3BT1 mutations is compared to the number of frames with the lowest RMSDs to mutant simulations excluded from the calculation of each ΔG value. The optimal parameter set of β = 0.002 and cutoff = 0.05% is used. Correlations are calculated after removing negligible predicted ΔΔG mutations. **(B)** For each mutation shown in Figure 3A, the backbone RMSD of each frame of the WT equilibrium 3BT1 dimer simulation to the average structure of that mutation’s equilibrium simulation is calculated. An average RMSD is then calculated for each mutation, and the distribution of these average values is shown for each set of mutations colored according to Figure 3A. **(C)** The correlation of predicted values to experiment is compared to total simulation time using the same set of mutations, parameters, and correlation calculation scheme as Figure 3A. The total simulation time is split evenly among three independent simulations, for example three independent 400 ns simulations are run for a total simulation time of 1200 ns. **(D)** The relationship between total simulation time and total number of frames with an RMSD to any mutant’s average structure of less than 1 Å is shown.

### MDR-FEP Relies on WT Equilibrium Simulations Sampling MUT-like Conformations

We initially predicted that the MDR-FEP method would detect mutational ΔG values by transiently sampling conformations which are more stable with mutant sidechains, and hence have more “low-RMSD frames” similar to conformations produced by mutant equilibrium simulations. We tested this hypothesis by evaluating the performance of the MDR-FEP method with differing numbers of low-RMSD frames removed, predicting that correlation to experiment would decrease as more low-RMSD frames are excluded. We indeed observed a strong relationship between correlation and the removal of low-RMSD frames, with overall correlation to experiment dropping from ∼0.55 to ∼0.30 with approximately 30% of the lowest-RMSD frames excluded (Figure 4A).

Likewise, we expected that for mutations whose predicted values disagreed with experiment, the WT equilibrium simulations would feature higher RMSDs to the MUT equilibrium simulations’ average conformations. We found that the WT equilibrium simulation was more likely to have higher RMSDs to the average conformation of mutations which were falsely predicted to be stabilizing to the 3BT1 dimerization process (Figure 4B). However, we did not observe this trend in the monomer simulations, perhaps indicating that both monomer and dimer simulations must sample low-energy frames to accurately predict ΔG (Figure S9).

Finally, we hypothesized that the WT equilibrium simulations would be more likely to sample low-energy frames with longer simulation times, thereby increasing correlation with experiment. We observed a strong relationship between correlation and total simulation time used (Figure 4C), as well as a direct relationship between total simulation time and total number of frames with sub-Angstrom RMSDs to mean structures from mutant equilibrium simulations (Figure 4D). These results indicate that the MDR-FEP method’s ability to predict experimental values depends on the WT equilibrium simulation’s transient sampling of low-energy mutant-like frames, and that this sampling depends on the total simulation time used. Based on these results, we believe that the performance of the MDR-FEP method on negligible predicted ΔΔG mutations, which suffer from insufficient sampling of low-energy frames, may improve with longer equilibrium simulation times.

### The MDR-FEP method selects conformations which are sterically favorable with the mutant but not WT sidechains

We hypothesized that the MDR-FEP method would be most sensitive to conformations which favor the mutant through a steric rather than electrostatic effect. In order to investigate this conformational selection mechanism, we analyzed the individual Rosetta score terms of sample long-distance mutations which were correctly identified to be destabilizing and stabilizing by the MDR-FEP method (Figure 5). We first analyzed the H143A mutation (Figure 5A-C), which is correctly predicted by the MDR- FEP method to destabilize 3BT1 dimerization (ΔΔGcalc = 4.8 REU, ΔΔGexp = 2.8 kJ mol^-1^). We found that excluding frames with ΔE < -500 REU resulted in a predicted ΔΔG not in accordance with experiment (0.7 REU). Analysis of the individual score terms of low-energy frames revealed high Lennard-Jones repulsion (“fa_rep”) between residues with the WT but not mutant sidechain, suggesting that mutating His143 to alanine in these particular conformations can relieve much more steric strain in monomeric state, resulting in destabilization of the dimer. A high correlation (R > 0.99) between the ΔE and Δfa_rep of individual frames in both the H143A monomer and dimer was observed.

**Figure 5.**
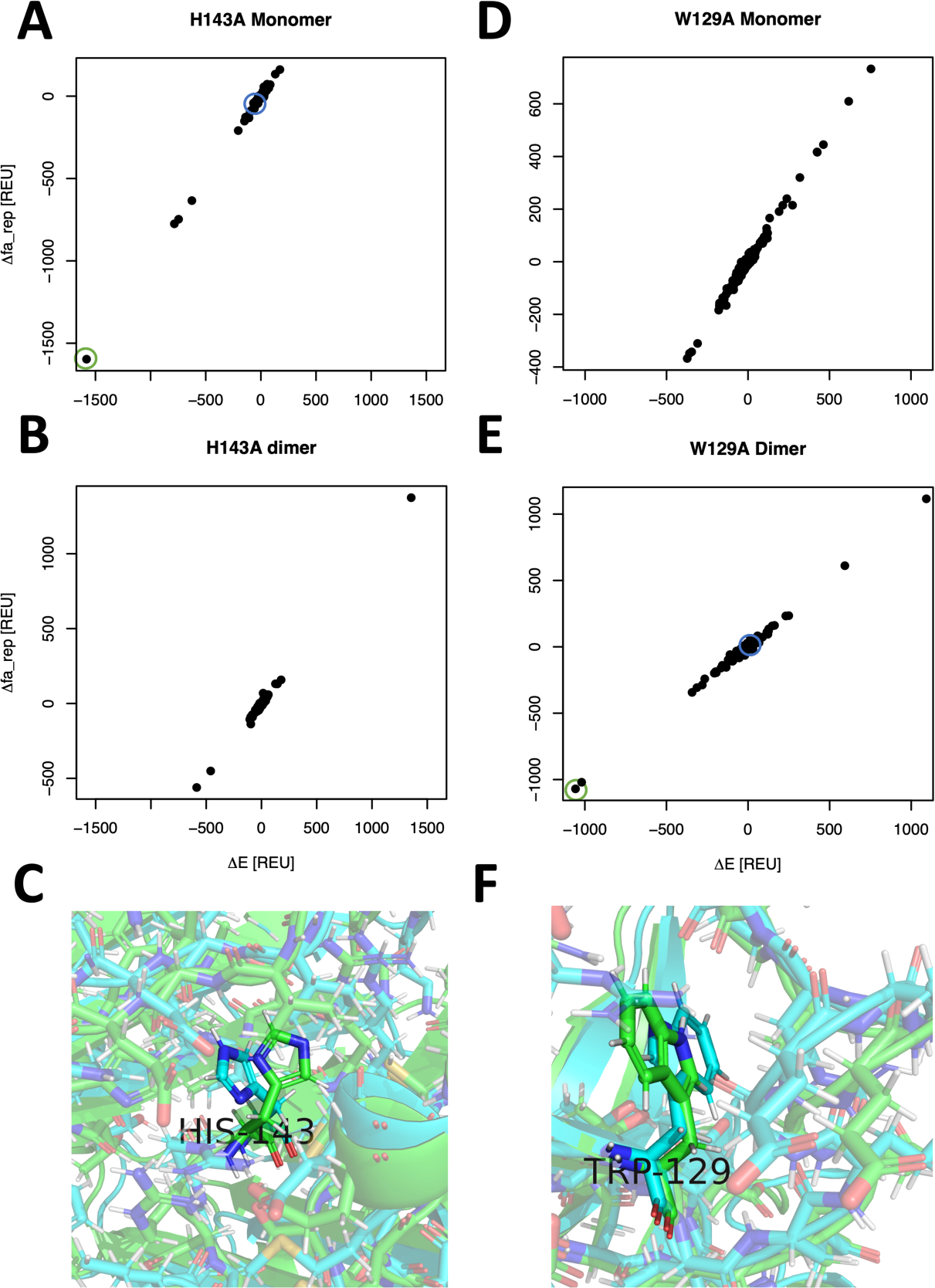
The MDR-FEP method selects conformations which sterically favor the mutant. **(A-B, D-E)** Plots comparing the overall ΔE and Δfa_rep (the Rosetta energy term which describes Lennard-Jones overlap) are shown for sample (A-B) destabilizing and (D-E) stabilizing mutations using the optimal parameter set of β = 0.002 and cutoff = 0.05%. Each frame is represented by a dot. **(C/F)** Protein images of the WT frames indicated by green circles in the (A)/(E) (above) are shown in green. Sample structures from frames with ΔE closer to 0, indicated by the blue circles in (A)/(E), are shown in blue for comparison.

Analysis of the W129A mutation, which was correctly predicted to stabilize dimerization, revealed similar trends (Figure 5D-F). When frames with ΔE < -500 REU were included, the MDR-FEP method correctly predicts the W129A mutation to stabilize dimerization (ΔΔGcalc = -2.9 REU, ΔGexp = -1.8 kJ mol^-1^). Removal of these frames resulted in an incorrect ΔΔG prediction of 1.7 REU. Score term analysis of these frames again revealed high steric overlap with the WT but not mutant sidechains present, indicating that mutation to alanine stabilizes the dimer by relieving more steric strain in the dimer than the monomer. The same strong relationship (R > 0.99) between Δfa_rep and overall ΔE was observed.

For the H143A mutation, we found that there were more frames with a calculated ΔE < -500 REU in the monomer than in the dimer. This higher number of low-energy frames contributes to a lower ΔG value in the monomer than in the dimer, resulting in an overall positive ΔΔG. Similarly, there were more low-energy frames in the dimer for W129A than in the monomer, resulting in a negative ΔΔG. Based on this analysis, we believe that the MDR-FEP method is able to correctly predict the ΔΔG values of these mutations by appropriately reproducing the relative number of mutation-favoring conformations in the monomer and dimer.

## Conclusions

Prediction of the effects of long-distance mutations remains an important goal for advancing the field of protein analysis and design. However, current methods which can accurately predict these long-range effects are limited by prohibitively large computational cost and are therefore not useful for large-scale analysis or protein design. Recent advancements using the Rosetta macromolecular modeling suite have allowed for accurate large-scale prediction of mutational ΔΔG values, however these methods are limited to interface mutations and cannot accurately detect long-distance effects. In the current study, we propose a method to combine the theoretical framework and extensive conformational sampling of alchemical free energy calculations with the high-throughput capabilities of Rosetta. In this approach, conformations from MD simulations are repacked and scored with Rosetta, and the effects of mutation- accommodating conformations are accounted for using theory based in statistical mechanics to allow for the accurate prediction of allosteric effects using only simulations of the WT system.

We evaluated the MDR-FEP method over three sets of mutations from two separate systems with accurate experimental data available. We found that the method is able to predict the effects of interface mutations with levels of accuracy comparable to current methods. Importantly, the MDR-FEP method was also able to predict the effects of long-distance mutations with similarly high levels of accuracy. We believe that this method represents a new approach to high-throughput analysis of mutations which propagate their effects over long distances.

While this work has focused on the bound and unbound states of two protein chains, the methodology can be generalized to any system where two alternate states can be defined. Other future applications to be explored include shifting the energy landscape of a protein towards particular conformational states, optimizing binding affinity/specificity for different small-molecules, or even stabilizing a small-molecule ligand/substrate in functionally active conformations.

## Materials and Methods

### Molecular Dynamics Simulations

Molecular dynamics simulations were generated *in silico* using the reference PDB structure 3BT1 (chains A and U) for the urokinase-urokinase receptor dimer, or 1BRS for the barnase-barstar dimer. Missing residues in 3BT1 were added using RosettaRemodel(39). All mutant 3BT1 structures were generated using the PyMOL software package(40). For all simulations, each PDB structure was placed in a dodecahedral box with 1.5 nm between the protein and box walls and solvated using the TIP3P water model(41). Joung Na^+^ and Cl^-^ ions were added to the simulation box at physiological concentrations of 150 mM(42). Each system was allowed to energy- minimize for up to 10,000 steps using the Steepest-Descent algorithm in GROMACS. Energy-minimized structures were then equilibrated with 20 ps of NVT, and then 20 ps of NPT, simulation using 1000 kJ mol^-1^ nm^-2^ all-atom position restraints. Each system was then equilibrated with three consecutive 20 ps NPT simulations using all-atom position restraints of 500, 250, and 125 kJ mol^-1^ nm^-2^, respectively. Finally, two successive 20 ps NPT equilibrations were run with force constants of 125 kJ mol^-1^ nm^-2^, first on all backbone atoms and finally on Cα atoms only.

Equilibrium simulations were run at constant temperature and pressure. The C-rescale barostat was used to hold pressure at 1 Bar, and the V-rescale thermostat used to hold temperature at 300 K. All system preparations were carried out using GROMACS functions and in-house scripts. Production simulation trajectories were generated using GROMACS 2021(43) with a 2 fs timestep, the Verlet neighbor-searching cutoff-scheme, and Particle-Mesh Ewald (PME) for van der Waals and electrostatic interactions. The AMBER 99SB*-ILDN(44, 45) force field was used to generate production trajectories, constraining all bonds using the LINCS algorithm(46).

### Protocol Implementation and Rosetta Repacking

For each system, three independent 500 ns simulations of both the monomer and dimer were run. Conformations were extracted every 1 ns from these trajectories. Each of these conformational frames was then repacked using the Rosetta 2020 fixed-backbone repacker, allowing for extra sub-rotamers for χ1 and χ2 angles. The “multi- cool-annealer” repacking option was used, where multiple “cooling” cycles are run and low temperature rotamer substitutions are then run from the 10 best network states generated during the cooling stage. Additionally, sidechain dihedral angle minimization was used on the lowest energy structure from the packer. For each frame, the score (*E*) from the lowest energy conformation of 50 repacking iterations was recorded with both the WT and mutant sidechains present. The following flags were used: `-ex1 -ex2 - multi_cool_annealer 10 -minimize_sidechains -ndruns 50`. Only residues which contained at least 1 atom within 10 Å of the mutation of interest in the crystal structure were repacked. For each mutation, the same set of residues was repacked in the monomer and dimer forms of the protein.

Following repacking and scoring, *ΔE* values for each frame were determined using the following equation:

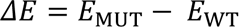

Where *ΔE* represents the difference in score between the frame with a WT and MUT sidechain present. The set of ΔE values generated for each mutation in the monomer and dimer states were converted into a distribution using kernel density estimation with a bandwidth manually set to 0.1, and a cutoff parameter was applied such that all values below the cutoff are set equal to zero. ΔG values were then generated from each distribution of *ΔE* values using the following equation:

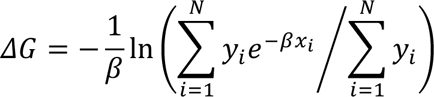

Whereand ; are the vectors describing the X and Y values of the kernel density function, respectively. Finally, ΔΔG values were generated using the following equation:

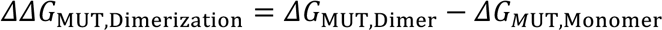

All experimental data in Figure 2 and Figure 3 were obtained from the SKEMPI 2.0 database(47) with mutations involving glycine removed, as were the locations of all mutations relative to the protein interaction interface (“interior” and “interface”). Flex ddG values shown in Figure 2 were calculated using the recommended parameters: `nstruct = 35, max_minimization_iter = 5000, abs_score_convergence_thresh = 1.0,` and `number_backrub_trials = 35000`.

### Data Analysis and Visualization

The solid lines in the data distributions shown in Figure 4B, Figure S8, and Figure S9 represent the mean value over three independent MD simulations. The shrouds in these distributions represent the standard error of these means, as calculated by

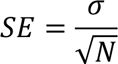

where σ is the standard deviation *N* is the number of independent trials. All protein images were generated using PyMol(40).

## Acknowledgements

This work was supported by NSF MCB2144396 (CAS). The authors would also like to thank Wesleyan University for computer time supported by the NSF under grant number CNS-0619508 and CNS-0959856, and the Extreme Science and Engineering Discovery Environment (XSEDE), which is supported by National Science Foundation grant number ACI-1548562 (under the allocation ID MCB190110), for resources on the SDSC Expanse and PSC Bridges compute clusters used to produce this work(48). To the best of our knowledge, no authors have any conflict of interest, financial or otherwise.

