## Supporting Figures for "Predicting binding affinity changes from long-distance mutations using MD simulations and Rosetta"

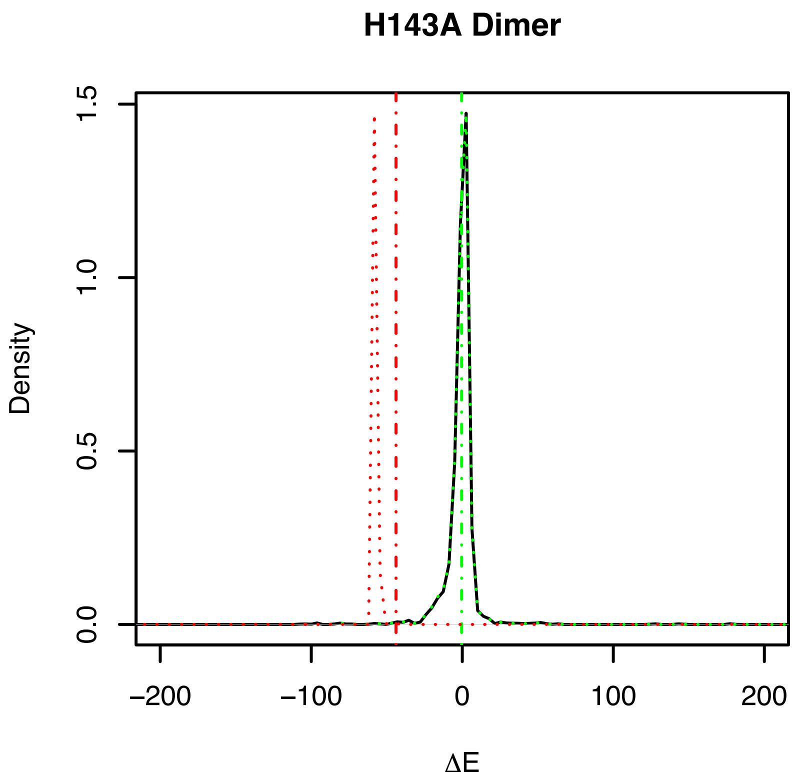


**Figure S1. High β values cause low-energy frames to dominate the ΔG calculation.**

The calculated ΔG values are shown as dot-dashed vertical lines, and the contribution of each portion of the ΔE distribution to the final ΔG value is shown as a dotted line. With the optimal β value of 0.002 (green), low-energy frames contribute appropriately to the overall ΔG value. Increasing β to 0.5 causes low-energy frames to dominate, resulting in inaccurate predicted ΔG values.

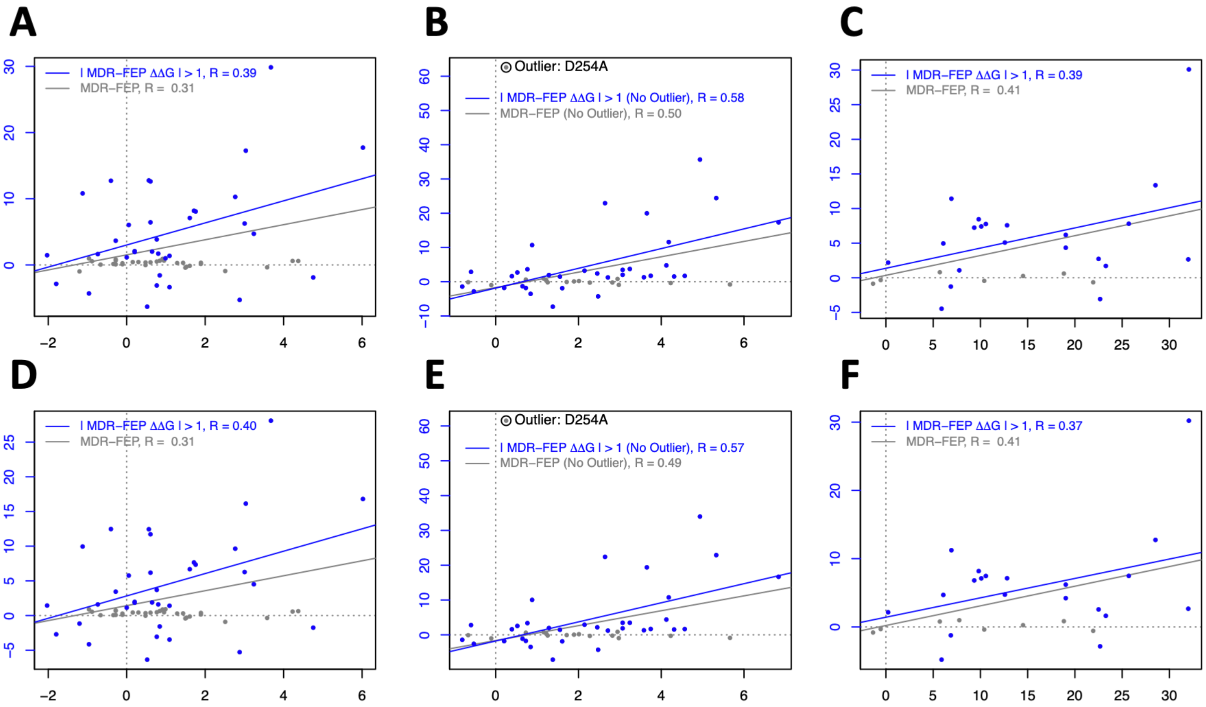


**Figure S2. MDR-FEP performance without the use of KDE.**

MDR-FEP correlation with experiment without the use of KDE is shown for (A) 3BT1 long-range interior, (B) 3BT1 interface, and (C) 1BRS interface mutations. The same datasets generated using KDE are shown below for comparison. Correlations are calculated using β = 0.002 with no cutoff applied. All correlations shown are statistically significant ($p > 0.05$).

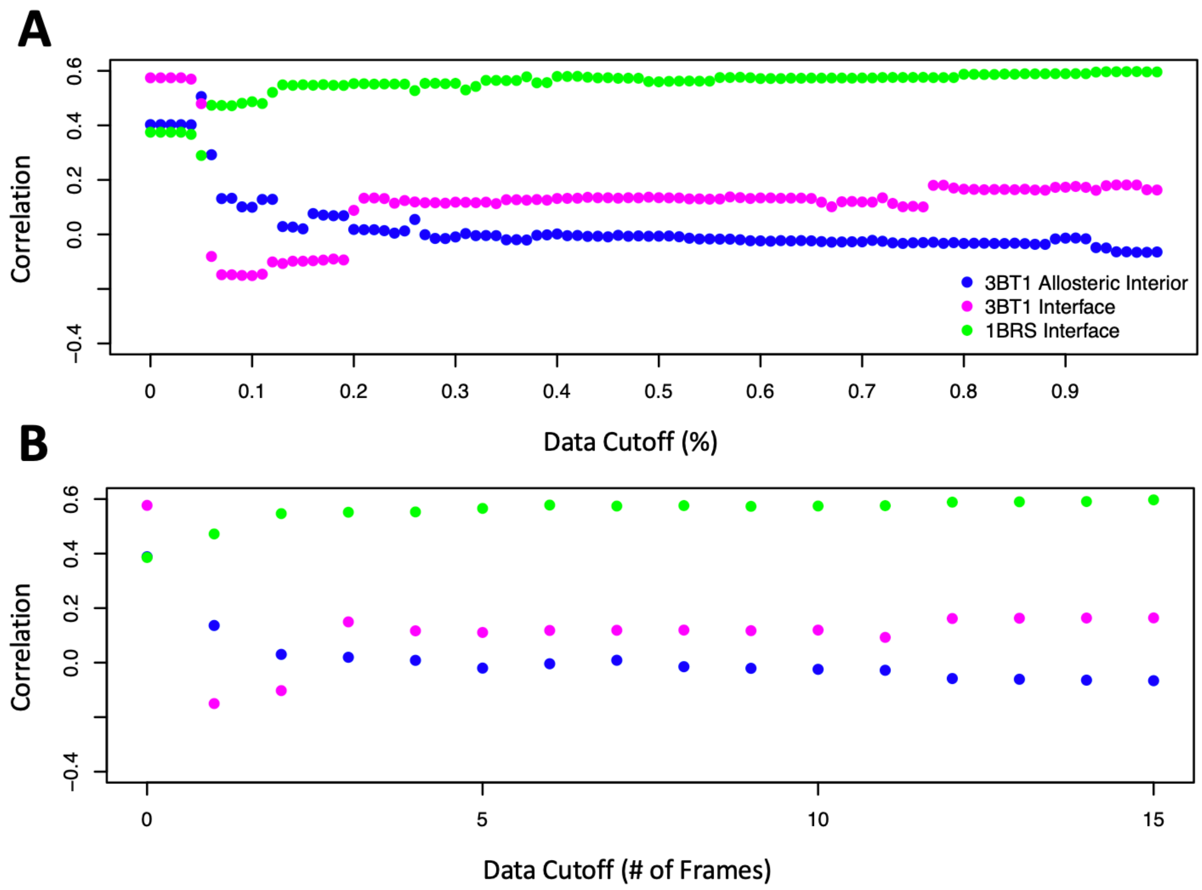


**Figure S3. The use of KDE allows for more precise data cutoffs to be applied.**

Correlation is compared to (A) a continuous cutoff percentage applied to the density of the data and (B) a discrete whole-frame cutoff applied to the raw data.

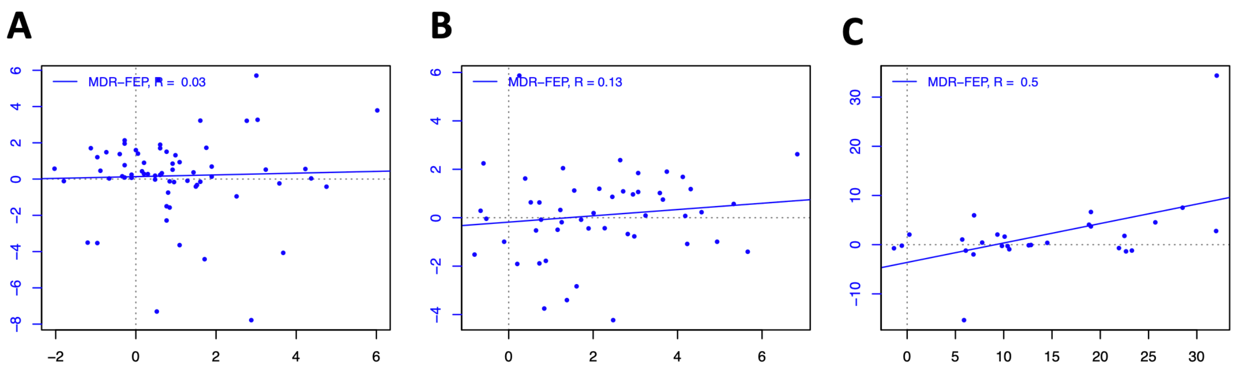


**Figure S4. MDR-FEP correlation to experiment using simple rather than exponential averaging.**

MDR-FEP correlation with experiment using simple averaging of ΔE distributions rather than exponential averaging is shown for (A) 3BT1 long-range interior, (B) 3BT1 interface, and (C) 1BRS interface mutations. Correlations are calculated using β = 0.002 with no cutoff applied.

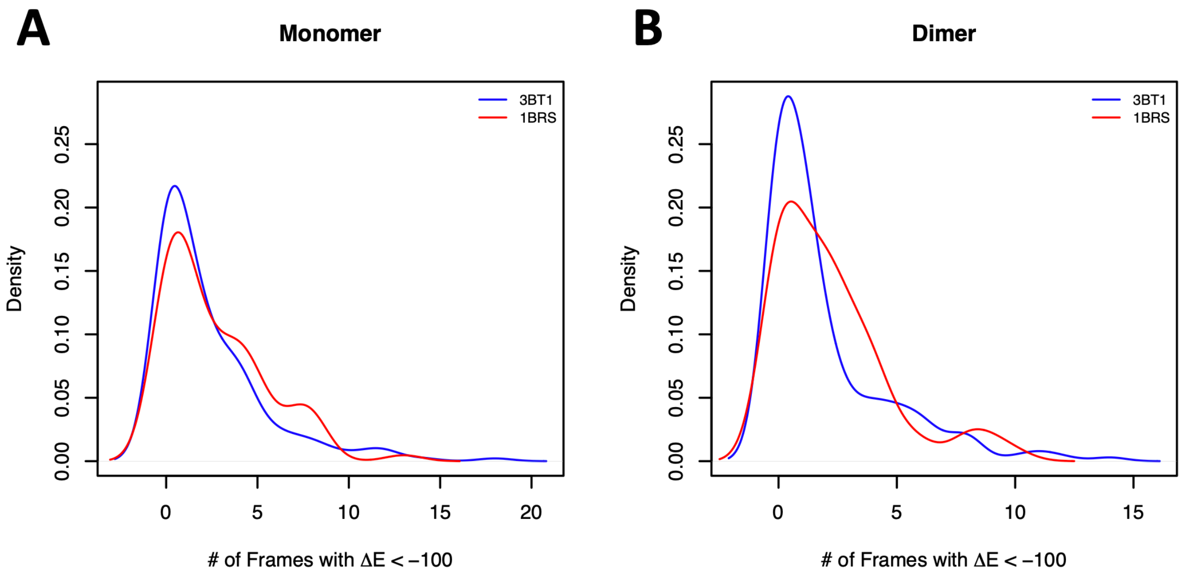


**Figure S5. 3BT1 and 1BRS simulations sample approximately equivalent numbers of low-energy frames.**

Comparison between the number of frames with an MDR-FEP predicted ΔE of less than -100 REU for mutations to 3BT1 and 1BRS in WT equilibrium (A) monomer and (B) dimer simulations.

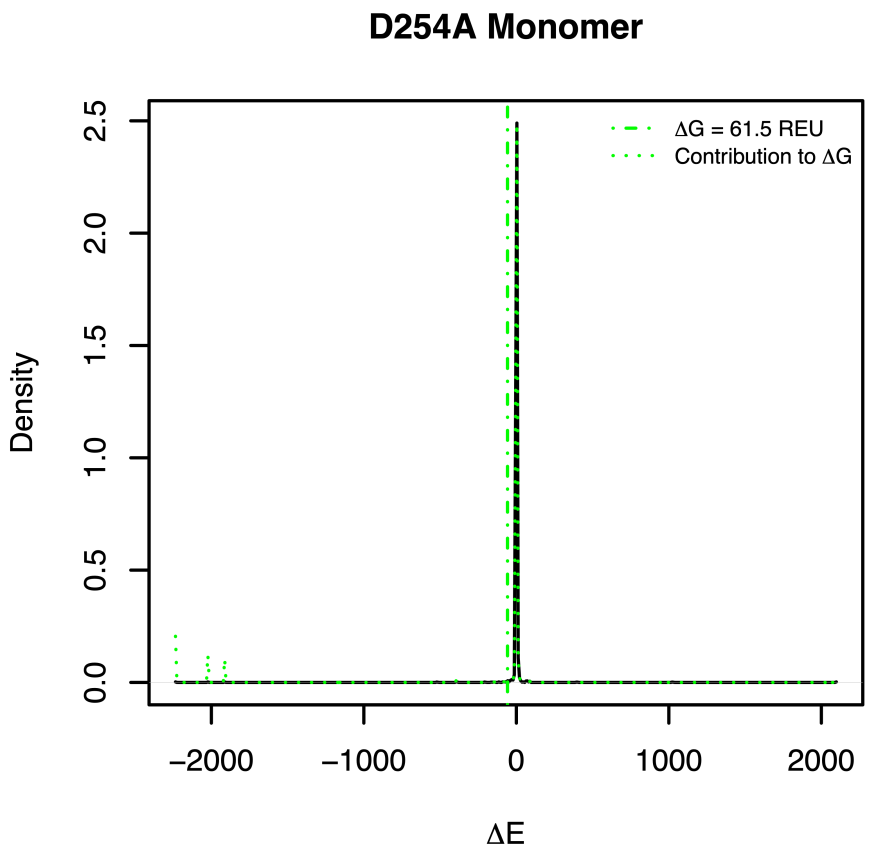


**Figure S6. The MDR-FEP method calculates an inappropriately large negative ΔG value for the D254A mutation, even at low β values.**

The ΔG value is calculated using the optimal parameter set of β = 0.002.0 and cutoff = 0.05. The fraction of the contribution (green dotted line) which falls within 99.9% of the main distribution (solid black line) is calculated to be 0.88.

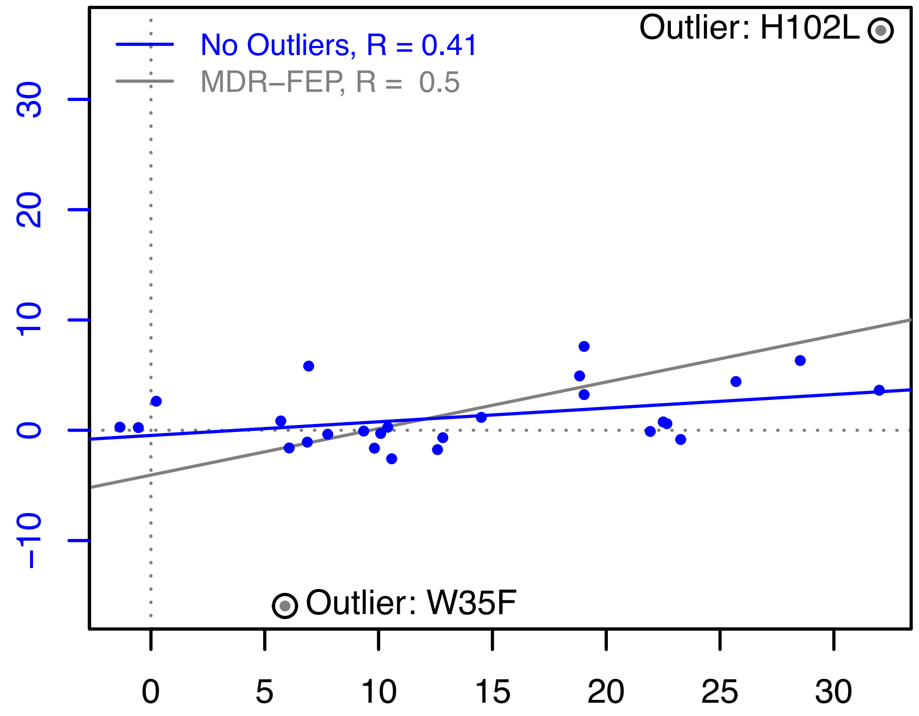


**Figure S7. MDR-FEP performance using the 1BRS dataset, excluding potential outliers. Correlations are calculated using the same parameter set as in Figure 2F.**

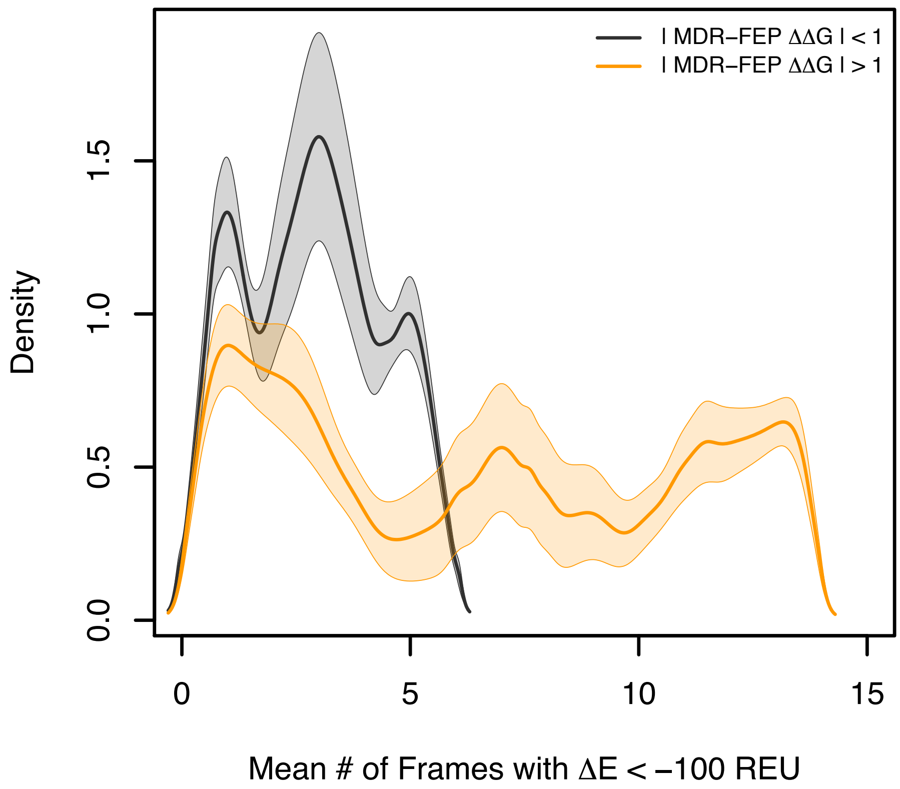


**Figure S8. The WT equilibrium simulation tends to sample fewer low-energy frames for negligible predicted ΔΔG mutations (gray) than mutations which have high predicted values (orange).**

Mutation set, parameters used, and coloring scheme are based on Figure 3A/D.

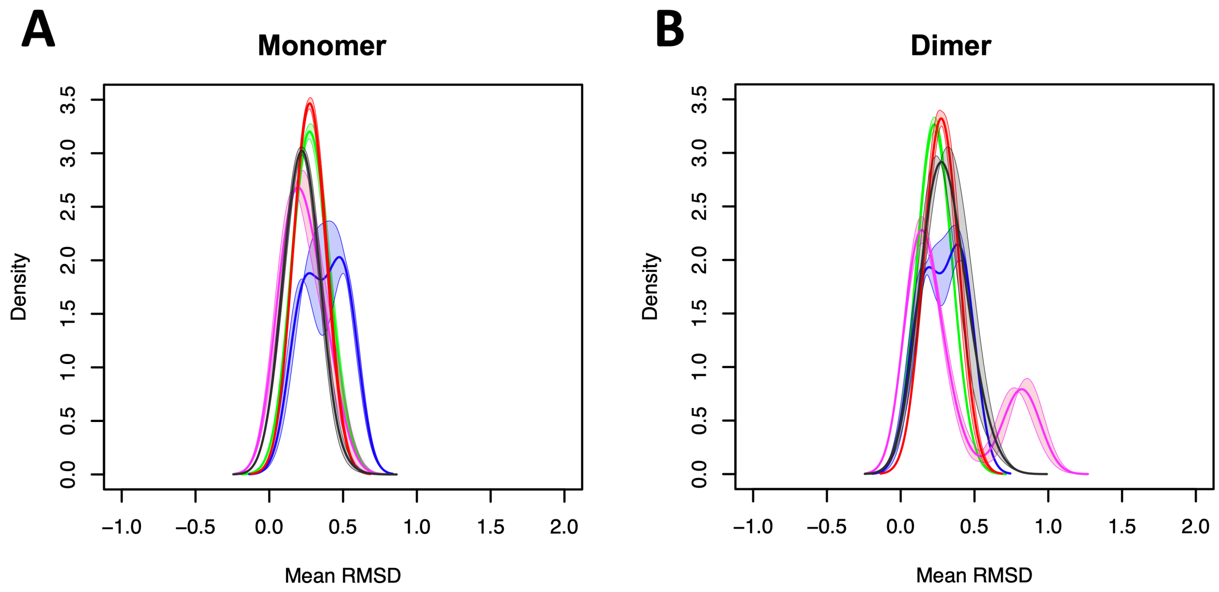


**Figure S9. WT equilibrium 3BT1 simulations looked less like mutant simulations for mutations which were incorrectly predicted to be stabilizing.**

The mean RMSD of the WT equilibrium 3BT1 simulations to the average structure generated by mutant equilibrium simulation is plotted for the sets of mutations described by the coloring of Figure 3 A-C.
